# Coordinated governance redirects agricultural expansion away from native vegetation in Brazil

**DOI:** 10.64898/2026.05.25.727617

**Authors:** Mario Barroso Ramos Neto, Alessandra Bertassoni, Marisa de O. Novaes, Giovanni Mallmann, Arley Haley Faria, Rafaela Albuquerque, Guilherme Ramos Vaz, Laerte Guimarães Ferreira

## Abstract

Tropical agricultural frontiers continue to drive native vegetation loss while supplying global commodity demand; yet it remains unclear how alternative governance pathways reshape where expansion occurs across interconnected biomes. Here, we use a spatially explicit land-use model to evaluate how contrasting governance regimes, with and without moderate livestock intensification, affect land-use futures across the Brazilian Amazon, Cerrado, and Pantanal through 2030. We compare Governance Inertia, Collaborative Governance, and Integrated Governance to assess their effects on native vegetation conversion, the land-use origins of soybean expansion, and landscape structure. Under scenarios in which conversion from native vegetation is explicitly constrained, projected soybean expansion remains nearly constant (∼6.2 Mha) but is redirected away from native vegetation and toward already converted lands. Relative to Governance Inertia, constrained-governance scenarios avoid approximately 13.5 Mha of native vegetation loss and require an estimated 13.9% increase in livestock occupation rate to maintain production with a smaller pasture footprint. These shifts are also associated with modest but consistent improvements in vegetation continuity and reduced fragmentation. Our findings show that the effectiveness of zero-deforestation trajectories depends not only on limiting expansion, but on governing where land-use demand is absorbed across tropical frontier systems.

## Introduction

Tropical regions account for a disproportionate share of both agricultural expansion and native vegetation loss. Across these frontiers, land-use change is increasingly shaped by the interaction between global commodity demand^1,2^ and regionally structured production systems^3^, making agricultural expansion not only a driver of biophysical transformation, but also a spatial governance challenge^4^. In South America, the coupled expansion of soybean production and cattle ranching has advanced across some of the world’s most biodiverse landscapes, often through frontier dynamics that extend beyond individual biomes and displace land-use pressure across connected territorial systems^5,6^. In this context, Brazil represents a critical case: it is simultaneously one of the world’s leading exporters of soybeans and beef^7^ and home to globally important reservoirs of biodiversity^8,9^. This dual role makes the country central to current debates on whether agricultural production and native vegetation conservation can be reconciled across tropical frontiers.

Brazil is also a particularly revealing case because strong formal governance coexists with uneven implementation. The country combines one of the world’s most comprehensive legal architectures for native vegetation protection, including the Forest Code, protected areas, and property-level environmental registration^10^, with persistent implementation gaps, unresolved liabilities, and recurrent political shifts in enforcement capacity^11^. As a result, governance in Brazil is not well described as simply present or absent. Its effects vary over time and across space, reflecting differences in legal obligations, monitoring capacity, land tenure conditions, and the institutional ability to enforce restrictions on land conversion^10,12,13^. These asymmetries are especially relevant across the Amazon, Cerrado, and Pantanal, where agricultural frontier dynamics are interconnected, but the governance conditions shaping land-use allocation are not uniform^12,14,15^.

Although recent advances in land-use modeling have improved the ability to project tropical deforestation and agricultural expansion under alternative socioeconomic and policy conditions^7,14,16–18^, important gaps remain. Many scenario-based assessments still focus primarily on aggregate land conversion, with less attention to whether governance changes ensure that expansion is absorbed within the land system ^14,18,14,18–20^. Yet this distinction is critical. In fact, ecological outcomes depend not only on how much native vegetation is lost, but also on how expansion is redistributed across land-use classes and how the remaining vegetation is spatially organized^21,22^. And there are few studies explicitly comparing governance counterfactuals within a consistent spatial framework while also examining the interactions among governance constraints, pasture-based intensification, and landscape structure across multiple biomes linked by agricultural frontier dynamics.

This gap is particularly important in frontier regions where soybean expansion, cattle production, and native vegetation conversion are tightly coupled^12,14,15^. In such settings, governance does not operate only by limiting total expansion; it also shapes the spatial pathways through which land-use demand is accommodated. Blocking or penalizing conversion from native vegetation may reduce direct pressure on natural areas, but it also creates adjustment pressures elsewhere in the land system, especially on already converted lands, such as pasture^1,23^. Understanding whether those adjustments can be absorbed without substantial changes in total agricultural expansion, and with minimal consequences for landscape structure, is therefore central to evaluating the practical meaning of zero-deforestation and deforestation-and conversion-free trajectories.

Here, we use a spatially explicit modeling framework to evaluate how alternative governance regimes, with and without moderate livestock intensification, reshape land-use futures across the Amazon, Cerrado, and Pantanal through 2030. We compare Governance Inertia, Collaborative Governance, and Integrated Governance to evaluate their effects on native vegetation conversion, the land-use origins of soybean expansion, and landscape structure. Because these governance pathways are implemented through explicit restrictions on transition pathways and spatial allocation, the central question is not whether native vegetation conversion declines under stronger governance, but how land-use demand is redistributed once those constraints are imposed. In doing so, we show how coordinated governance can redirect agricultural expansion toward already converted lands, while improving indicators of landscape functional integrity across interconnected tropical frontiers.

## Results

The three governance pathways produced similar totals of projected soybean expansion by 2030, but differed in how land-use change was allocated across land-use classes and in their implications for native vegetation retention and landscape structure (see Supplementary Information for results of all scenarios). Because the Collaborative Governance (CG) and Integrated Governance (IG) scenarios explicitly modify transition pathways and constraints relative to

Governance Inertia (BAU), lower native vegetation conversion under these regimes should be interpreted as the expected consequence of the imposed counterfactual rules. The key analytical question, therefore, is how land-use demand is reallocated once these additional constraints are introduced.

### Governance pathways reduce native vegetation conversion by design

Relative to BAU, both CG and IG projected substantially lower native vegetation conversion between 2023 and 2030. Under BAU, pasture expanded by 7.03 Mha over the period, whereas pasture area declined by approximately 7.53 Mha under both CG and IG, corresponding to a reduced pasture occupation of about 14.5 Mha relative to BAU (Fig. 1B; Figure S2). In turn, both constrained-governance scenarios avoided approximately 13.52 Mha of native vegetation loss compared with BAU. Because these scenarios explicitly constrain major pathways of native vegetation conversion, the resulting reduction in vegetation loss reflects the imposed governance assumptions embedded in the scenario design. Indeed, the more informative result is that, once those restrictions are imposed, the model reallocates expansion pressure away from native vegetation and toward already converted lands. Pasture expansion under BAU was strongly associated with native vegetation loss (Spearman ρ = 0.91; p < 0.001; Figure S3), reinforcing the central role of pasture-mediated conversion in the baseline configuration.

**Figure 1.** Governance regimes and their effects on land-use dynamics and native vegetation loss. (A) Conceptual representation of the three governance regimes: Business as Usual (BAU), Collaborative Governance (CG), and Integrated Governance (IG), highlighting their main mechanisms. (B) Spatial distribution of avoided native vegetation loss under IG relative to BAU (2023–2030), expressed as the percentage of unconverted native vegetation within each 20 km² hexagonal cell. Color classes indicate increasing levels of avoided loss. (C) Prior land-use composition of soybean expansion under each governance regime, showing the contribution of native vegetation, pasture, and other land uses to total expansion, along with the spatial distribution of expansion intensity. (D) Schematic representation of livestock intensification under IG, indicating a modest increase in stocking rate (from 1.00 to 1.14 AU ha).

### Governance redirects soybean expansion toward already converted lands

Projected soybean expansion remained nearly constant across the three scenarios, totaling approximately 6.2 Mha by 2030 (BAU: 6,262,152 ha; CG: 6,226,582 ha; IG: 6,226,789 ha). Thus, the governance scenarios did not substantially alter the total amount of soybean expansion. Instead, they changed its origin (Fig. 1C). Under BAU, 8.9% of soybean expansion originated from native vegetation (555,703 ha) and 12.1% from pasturelands (755,265 ha). Under CG and IG, the contribution of native vegetation declined to 7.6% (∼0.47 Mha), while pasture contributions were slightly reduced to 11.4–11.5% (∼0.71 Mha). Across all scenarios, most soybean expansion occurred over other pre-existing anthropogenic land uses, mainly temporary crops and mixed-use mosaics, which accounted for roughly 79–81% of total expansion (Figure S4). These results indicate that governance does not impact soybean expansion itself; rather, it redirects where that expansion is sourced from within the land-use system.

### Moderate intensification accommodates pasture contraction under constrained governance

The contraction of pasture area under the alternative governance regimes implies that equivalent livestock production would need to be maintained with a smaller pasture footprint. Based on the ratio between pasture area under BAU and under the constrained-governance scenarios, the estimated intensification requirement is 13.86% (Fig. 1D), equivalent to an increase in the average national stocking rate from 1.00 to approximately 1.14 AU ha⁻¹ (animal unit per hectare)^24^. Because CG and IG generate nearly identical reductions in pasture area, this requirement is also nearly identical in both scenarios. In this framework, moderate intensification should be interpreted as an accommodating mechanism that helps sustain production once native vegetation conversion is constrained, rather than as the primary explanation for lower native vegetation loss. This interpretation is consistent with the logic of the scenario design, in which the main conservation effect arises first from explicit restrictions on conversion pathways, while intensification affects how production is reorganized within the already converted land base.

### Governance-driven land reallocation improves structural landscape integrity

Differences in land allocation across scenarios were also reflected in landscape structure. Using the Landscape Line Intercept (LLI), we found a redistribution of landscape configurations under constrained governance relative to BAU (Fig. 2). Under BAU, 7.5% of our basic unit of analysis, i.e. 20 km^2^ hexagons, were classified as highly fragmented, with low continuity and low native vegetation area, whereas 53.1% were classified as intact landscapes, with high continuity and high native vegetation area (Fig. 2A). Under IG, the share of intact landscapes increased, and the share of highly fragmented landscapes declined (Fig. 2B). Relative to BAU, IG showed an increase of 1.7 percentage points in high-continuity, high-vegetation hexagons and a decrease of 2.7 percentage points in low-continuity, low-vegetation hexagons, with smaller changes in intermediate classes (Fig. 2C). Spatial patterns under IG closely resembled those observed under CG (Figure S5,6). Taken together, these results indicate that once native vegetation conversion is explicitly constrained and agricultural expansion is redirected towards already converted lands, the resulting land-use allocation is associated not only with greater native vegetation retention, but also with more continuous and less fragmented landscape configurations.

**Figure 2.** Changes in landscape structure under different governance regimes. (A) Relationship between native vegetation area (%) and landscape continuity (%) under the Business as Usual (BAU) scenario in 2030, with the corresponding distribution of areas across classes of vegetation cover and continuity. (B) Same relationship under the Integrated Governance (IG) scenario in 2030. (C) Difference between IG and BAU, showing changes in the proportion of areas across vegetation cover and continuity classes. Positive values (green) indicate an increase in more continuous landscapes, whereas negative values (red) indicate a shift toward more fragmented conditions. Axes represent classes of native vegetation area (%) and continuity (%), highlighting transitions between fragmented and continuous landscape configurations.

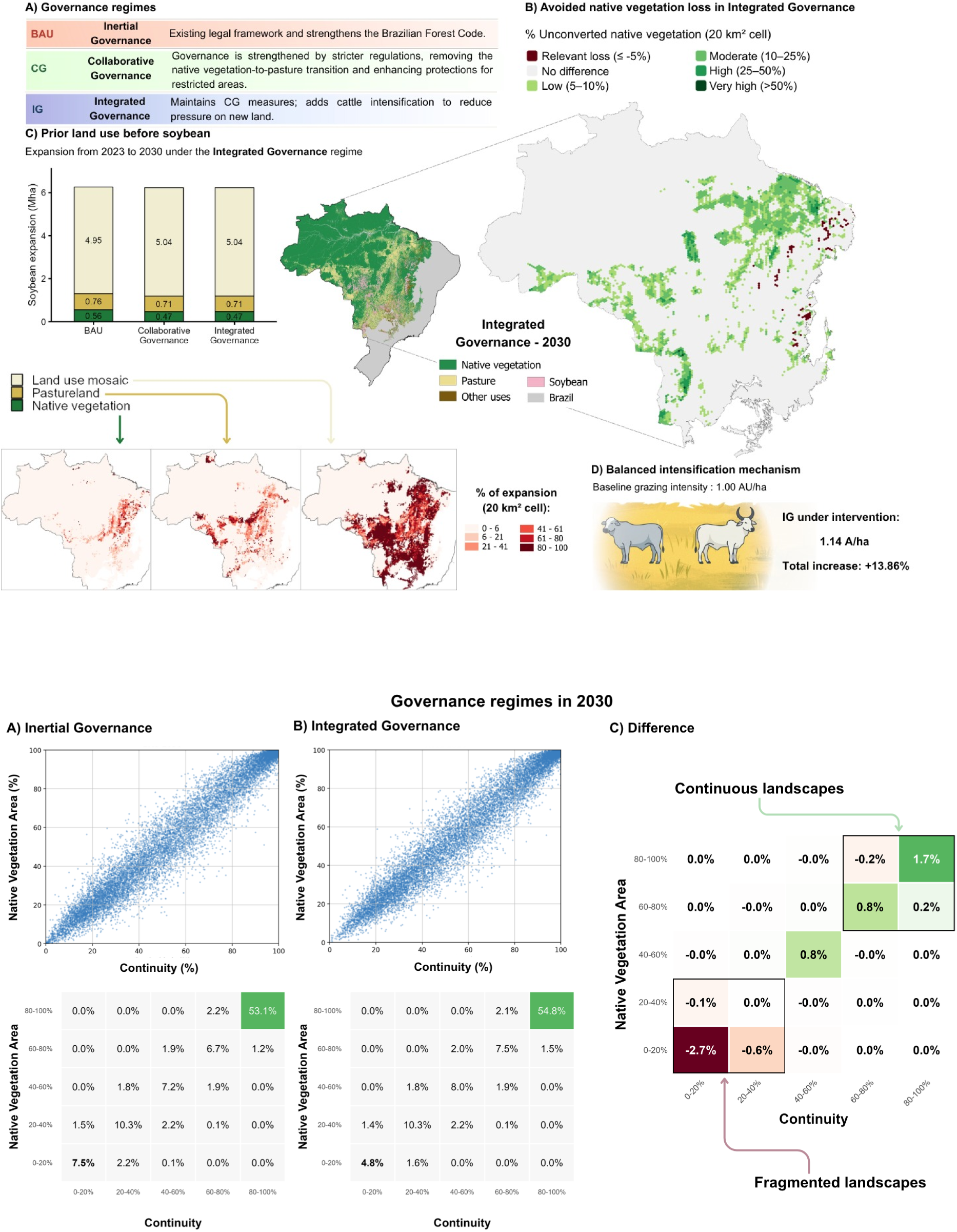

## Discussion

A first implication of our results is that public governance remains the institutional backbone of zero-deforestation trajectories in Brazil^10,11^. In our framework, the lower levels of native vegetation conversion under Collaborative Governance and Integrated Governance are the expected consequence of explicit counterfactual restrictions on key conversion pathways and stricter normative constraints in spatial allocation (relative to BAU). In other words, the more positive outcome is a consequence of reinforcing environmental rules and improved territorial occupation: soybean expansion remains broadly stable in total area, but is reallocated away from native vegetation and towards already converted lands. This distinction is important because it clarifies what this modeling exercise is designed to evaluate. Rather than showing that deforestation would decline spontaneously under stronger governance, the scenarios examine whether agricultural expansion can be spatially redirected once public and hybrid governance constraints are made effective.

This interpretation places Brazil’s Forest Code and the control of illegal deforestation at the center of the discussion. In practice, public governance in Brazil is grounded in a dense legal framework^10,11^, but its effects are neither temporally stable nor spatially uniform. They are highly sensitive to political continuity, enforcement capacity, and the administrative priorities of each federal cycle^11,25^. This sensitivity is not only institutional, but territorial. Under the Forest Code, Legal Reserve requirements are asymmetric by design: 80% in forest areas within the Legal Amazon, 35% in Cerrado areas within the Legal Amazon, and 20% in the rest of the country (Law 12,651/2012). As a result, similar improvements in monitoring or enforcement are unlikely to produce equivalent outcomes across the Amazon, Cerrado, and Pantanal. In this sense, stronger governance should not be understood as a homogeneous condition across the study area, but as a spatially differentiated combination of legal obligations, enforcement intensity, and implementation capacity. Brazil’s recent policy trajectory reinforces this point: federal plans to mitigate deforestation in the Amazon and Cerrado were discontinued in 2020 and resumed in 2023, underscoring how governance gains and setbacks remain tied to political cycles^26–28^.

Within this broader public framework, private and hybrid governance mechanisms matter most when they reinforce, rather than replace state regulation^29^. Our results suggest that this complementarity is especially relevant in interconnected agricultural frontiers^30^, where the central challenge is not simply whether expansion occurs, but where it is redirected under constraint. In that sense, zero-deforestation and conversion-free initiatives are likely to be most effective when they operate as spatial coordination mechanisms layered on top of an existing public regulatory framework^19,31,32^. This is also why the differences between BAU and the constrained-governance scenarios are more informative than the relatively small differences between CG and IG in avoided native vegetation conversion. Once a major conversion pathway from native vegetation is blocked, the main analytical question becomes how the remaining land-use demand is reorganized across the already converted land base. That logic also helps explain why soybean expansion remains nearly constant across scenarios, even as its land-use origin changes.

The role of livestock intensification should be interpreted within that same sequence. In our simulations, moderate intensification does not, by itself, create the conservation effect; rather, it serves as an accommodating mechanism once conversion constraints are already in place. By allowing equivalent livestock production to be maintained with a smaller pasture footprint^33,34^, intensification helps absorb part of the adjustment required by governance-driven pasture contraction.

This interpretation is more defensible than presenting intensification as a universal solution to frontier expansion. Literature has long emphasized that productivity gains can either spare land or stimulate renewed expansion, depending on the surrounding institutional context^4,31^. Our results support the former interpretation only under coordinated governance conditions^29,35^, where restrictions on conversion and spatial reallocation towards already converted lands operate simultaneously. In this sense, intensification is best understood as a mechanism that helps stabilize production under stricter land-use constraints, rather than as an independently sufficient route to zero deforestation^36^.

The governance effects extend beyond aggregate land conversion and become visible in the structural organization of landscapes^37^. Once agricultural expansion is redirected away from native vegetation and towards previously converted lands, the resulting land-use allocation is associated with more continuous vegetation configurations and fewer highly fragmented landscapes. This does not mean that the observed shifts are large in absolute terms, but their direction is consistent and ecologically meaningful. Spatial arrangement matters because biodiversity persistence, microclimatic stability, disturbance propagation, and movement across heterogeneous landscapes depend not only on habitat amount, but also on how remnant vegetation is configured^21,38^. Here, the LLI is therefore most useful as an indicator of structural landscape integrity: it supports the argument that governance affects not only how much vegetation is retained, but also how remaining vegetation is spatially organized.

Another implication is that governance pathways of the kind represented here are increasingly tied to financial mechanisms that can turn legal and voluntary commitments into operational land-use transitions^39,40^. This is no longer a hypothetical policy space in Brazil. Public programs such as the ABC+ Plan (www.gov.br/agricultura/pt-br/assuntos/sustentabilidade/planoabc-abcmais) and the National Program for the Conversion of Degraded Pastures (Federal Decree 11.815/2023) explicitly promote the recovery and productive reuse of degraded pasturelands as a route to expand production without further pressure on native vegetation. More recent instruments point in the same direction. The Eco Invest program (www.gov.br/tesouronacional/pt-br/fomento-ao-investimento) has been structured to mobilize capital for large-scale recovery of degraded lands, while the Responsible Commodities Facility now has international climate-finance backing to support deforestation- and conversion-free soy production in the Cerrado. At the same time, payment-for-environmental-services schemes such as Floresta+ Amazônia (www.florestamaisamazonia.org.br/) demonstrate that direct conservation incentives are also becoming part of the implementation landscape. Together, these examples suggest that the feasibility of DCF governance and pasture productivity gains increasingly depend on finance, credit, and risk-sharing arrangements that reduce the cost of compliance and make land-use reallocation economically viable. These findings also help clarify what this study can and cannot say about leakage and indirect land-use change. In the constrained-governance scenarios, soybean expansion does not simply disappear; it is redistributed. That result is encouraging because it suggests that, under explicit restrictions on native vegetation conversion, a substantial share of expansion can be accommodated on already converted land while maintaining overall soybean growth. At the same time, the model does not imply that displacement risks vanish under real-world conditions. Whether reallocation reduces or amplifies indirect deforestation depends on actor behavior, market incentives, enforcement quality, and the availability of suitable converted land in the territory^41^. For this reason, our results should be interpreted as evidence that coordinated governance can improve the spatial allocation of expansion, not as a statement that indirect effects would be fully contained in practice.

Several caveats should therefore be made explicit. First, because governance scenarios are implemented through imposed restrictions on transition pathways and normative weights, and the reduction in native vegetation conversion is a designed counterfactual consequence of the scenario architecture. Thus, the more informative result lies in the subsequent reallocation of land-use demand. Second, the model does not include several mechanisms that can drive deforestation in practice, even under formal legal constraints. These include clearing motivated by land speculation and asset valorization, occupation of public lands followed by attempts at regularization, strategic deforestation to consolidate tenure claims, infrastructure-led frontier opening, and rapid behavioral responses to commodity-price shocks. Such mechanisms may induce clearings that are only partially related to immediate productive demand, thereby weakening the correspondence between productivity gains, legal restrictions, and actual land-use outcomes. Third, the model does not explicitly represent political discontinuities in enforcement, selective compliance by heterogeneous actors, or the expectation of future amnesty, all of which can alter the effectiveness of public governance over time. Finally, the landscape metric used here captures structural continuity rather than realized functional connectivity and should thus be interpreted as an indicator of fragmentation-related landscape organization rather than a direct measure of ecological movement or demographic persistence.

Taken together, our results suggest that the core challenge of tropical land-use governance is not simply to reduce expansion in the aggregate, but to determine where and under which constraints expansion is absorbed across interconnected territorial systems. In Brazil, that challenge is anchored first in the public governance architecture of the Forest Code and illegal-deforestation prevention, but it increasingly depends on complementary private commitments and financial mechanisms that make compliance and land-use reallocation feasible at scale. Under those conditions, moderate livestock intensification can help maintain production while reducing dependence on new land clearing, and the resulting redistribution of expansion can generate not only avoided native vegetation conversion but also improvements in structural landscape integrity. The broader implication is that zero-deforestation trajectories may depend less on inventing entirely new regulatory frameworks than on making existing rules more spatially effective, politically durable, and financially implementable across heterogeneous frontier regions. The future of tropical landscapes will depend not only on how much commodity expansion occurs, but also on how effectively governance redirects that expansion away from native vegetation and towards already converted or degraded lands.

## Methods

### Spatially explicit land-use modeling framework

We used a spatially explicit land-use and land-cover (LULC) model based on a stochastic cellular automata to simulate land-use dynamics across the Brazilian Amazon, Cerrado, and Pantanal through 2030. Simulations were conducted in Dinamica EGO (version 8)^42^, which was chosen for its ability to capture spatial dependence and neighborhood effects. The analysis was performed at 30 m spatial resolution and focused on four aggregated land-use classes: native vegetation, pasture, soybean, and other uses. The latter included other agricultural classes and mixed-use anthropogenic mosaics. This aggregation was adopted to retain the main transitions relevant to agricultural frontier dynamics while keeping the modeling framework parsimonious.

Model calibration used observed land-use transitions between 2018 and 2023, with the 2018 and 2023 MapBiomas Collection 9^43^ maps defining the initial landscape and the observed end state, respectively. Simulations were then carried out from 2023 to 2030. To represent spatial heterogeneity, we divided the study area into 24 spatially constrained subregions, based on data-driven regionalization of recent land-use composition and transition rates (Fig. S1). Calibration and simulation were conducted independently within each subregion, and outputs were mosaicked for final analysis. All raster layers were reprojected, resampled, and aligned to a common grid prior to modeling. Detailed descriptions of the preprocessing procedures, regionalization framework, transition matrix, and explanatory variables are provided in Supplementary Methods 1–3.

### Scenario design and implementation

We evaluated three counterfactual scenarios representing contrasting governance pathways: Business as Usual (BAU), Collaborative Governance (CG), and Integrated Governance (IG). Across all scenarios, historical transition rates estimated from the 2018–2023 period were used as the empirical basis for annual land-use change.

Governance effects were implemented by modifying spatial allocation mechanisms, rather than by changing aggregate transition rates. Accordingly, differences in projected native vegetation conversion between scenarios should be interpreted as the expected consequences of the respective imposed rules.

In BAU, observed transition dynamics were projected forward relatively to baseline governance conditions. Legal protection variables remained in the model as partial normative constraints, reflecting the presence of formal regulation under persistent implementation gaps. In CG, governance was strengthened through two structural changes: the removal of the native vegetation-to-pasture transition from the permitted transition matrix, and the assignment of stricter normative weights to protected and legally restricted areas. This scenario was designed to represent a governance regime in which public regulation is reinforced by private and hybrid arrangements that reduce the probability of expansion into native vegetation. In IG, the spatial containment rules of CG were preserved, and a livestock intensification suitability layer was added to redirect production pressure toward already converted pasturelands. This layer was derived from a degraded pasture mitigation framework and was used to favor the continued productive use of pasture areas rather than expansion into native vegetation^44,45^. These scenario settings should be interpreted as policy-relevant counterfactual parameters rather than as empirical estimates of realized enforcement intensity.

### Calibration and validation

Spatial allocation was calibrated using the Weights of Evidence (WoE) approach^46^, which estimates the statistical association between observed transitions and explanatory variables through Bayesian inference. Transition matrices were first calculated from the 2018 and 2023 LULC maps for each subregion. These matrices defined the amount of projected annual change, while WoE-derived probability surfaces determined where transitions would be preferentially allocated. To reduce redundancy in the allocation surfaces, highly correlated explanatory variables were screened before model fitting. Legal protection variables such as Legal Reserves, Permanent Preservation Areas, Strictly Protected Areas, and Indigenous Lands were retained as normative modifiers even when their empirical signal in the calibration period was weak, because the objective of the scenarios was to represent institutional counterfactuals rather than to rely exclusively on observed recent enforcement outcomes^47^.

Spatial allocation was implemented using Dinamica EGO’s Patcher and Expander functions, which regulate the balance between the formation of new patches and the expansion of existing ones^42^. Model performance was evaluated by comparing simulated 2023 maps with observed MapBiomas 2023 maps using neighborhood-based similarity metrics. Validation was conducted with moving windows from 3 × 3 to 11 × 11 pixels, and the 11 × 11 window was used as the main benchmark because it provided a balance between fine spatial precision and neighborhood representation at 30 m resolution. Regions were considered adequately simulated when they met a constant-decay similarity threshold of at least 0.4, or, when achieved, an exponential-decay similarity threshold above 0.5^42^. When these criteria were not initially met, patch-allocation parameters were recalibrated, and the model was re-evaluated. Full variable lists, summary table for WoE, correlation test results, allocation parameters, and regional validation outcomes are provided in Supplementary Methods 4-6.

### Post-simulation analyses

We quantified four outcome dimensions from the simulated 2030 landscapes. First, the avoided native vegetation conversion was calculated as the difference in projected native vegetation loss between BAU and each constrained-governance scenario. These contrasts were summarized for the full study area and spatially represented using a hexagonal grid to show geographic variation in avoided conversion. Second, the origin of soybean expansion was identified by comparing each simulated 2030 map with the 2023 baseline and recording the prior land-use class of each newly converted soybean pixel. This allowed soybean expansion to be partitioned among native vegetation, pasture, and other already converted land uses. Third, the livestock intensification required to accommodate the reduced pasture area under each alternative scenario was estimated as the ratio of total pasture area under BAU to that under the corresponding constrained-governance scenario. This ratio was then applied to a baseline national stocking rate of 1.00 animal unit per hectare to estimate the effective stocking rate needed to sustain equivalent livestock production with a smaller pasture footprint. Because the conservation effect in the scenarios is imposed first through governance constraints, this metric was used to evaluate the production adjustment required under those constraints rather than as an independent driver of avoided native vegetation conversion.

Fourth, we assessed structural landscape integrity using the Landscape Line Intercept (LLI). For each 20 km² hexagonal unit, LLI combines two structural dimensions: the proportion of native vegetation within the cell and the continuity of native vegetation along the cell perimeter. The first component captures habitat amount within the cell, whereas the second captures boundary continuity with the adjacent landscape context. Both components were calculated from binary native-vegetation rasters derived from the 2023 MapBiomas baseline and from each 2030 simulated scenario. In this manuscript, LLI is used as an indicator of landscape structure associated with fragmentation and vegetation continuity, rather than as a direct measure of functional connectivity. Expanded derivation and additional methodological details of the perimeter-based metric are provided separately in Supplementary Methods 7.

### Reproducibility and reporting

All processing was conducted in reproducible workflows, and code used for data processing and simulation is available in the project repository. To keep the main text focused on the logic of the policy experiment, additional details on raster preprocessing, regionalization, explanatory variables, WoE calibration, allocation parameters, scenario tables, and expanded LLI computation are provided in the Supplementary Information.

## Data Availability

All data supporting the findings of this study are available within the paper, its Supplementary Information files and in publicly accessible repositories under the https://github.com/lapig-ufg/Scenarios-project and https://zenodo.org/records/20159165.

Supplementary materials list:

Supplementary Methods 1. Input data and preprocessing

Supplementary Methods 2. Regionalization framework

Supplementary Methods 3. Transition matrix and explanatory variables

Supplementary Methods 4. Weights of Evidence calibration and spatial allocation

Supplementary Methods 5. Scenario parameterization

Supplementary Methods 6. Validation procedures

Supplementary Methods 7. Post-simulation calculations

Supplementary Results 1. Full three-scenario outputs

## Code availability

All code used for data processing and statistical analyses has been deposited in a publicly accessible GitHub repository under the https://github.com/lapig-ufg/Scenarios-project.

ChatGPT (OpenAI) was used to assist with language editing and clarification of the final manuscript, while the authors retain full responsibility for all scientific content, analysis, and conclusions.

## Author Contributions

M.B.R.N. conceived the study, which was designed by A.B., M.O.N., A.H., L.G.F., and M.B.R.N. M.O.N., A.H., G.M., R.A., and G.R.V. collected data. A.B., M.O.N., G.M., R.A., G.R.V., and M.B.R.N. performed the analysis. A.B., M.O.N., and M.B.R.N. wrote the first draft of the article. All the authors contributed to the discussions and paper revision. All authors read and approved the final manuscript.

