## Supplementary Information for "Coordinated governance redirects agricultural expansion away from native vegetation in Brazil"

[1]The Nature Conservancy (TNC)

[2]Instituto Nacional da Mata Atlântica (INMA)

[3]Remote Sensing and GIS Laboratory (LAPIG/UFG)

### Supplementary Methods 1. Input data and preprocessing

The data acquisition strategy aimed to capture fine-scale land-use transitions, prioritizing spatial granularity to represent neighborhood-level dynamics. Land-cover data were sourced from MapBiomas Collection 9 (https://brasil.mapbiomas.org/), a high-resolution initiative producing annual LULC maps derived from Landsat imagery. For this study, the 2018 map served as the baseline landscape and the 2023 map as the final landscape, defining the transition period used to estimate change rates.

The integration of data from the National Rural Environmental Registry System (SICAR) and the Land Management System (SIGEF) was essential for representing land tenure and institutional constraints. These datasets enabled property-level representation of Legal Reserves (LRs), Permanent Preservation Areas (PPAs), and the distinction between private and public lands. The Forest Code compliance layer incorporated biome-specific Legal Reserve thresholds defined by Brazilian Law 12.651/2012: 80% for forest formations in the Legal Amazon, 35% for Cerrado formations within the Legal Amazon, and 20% elsewhere.

All spatial data were harmonized to a common 30 m grid and projected to South America Albers Equal Area Conic (ESRI:102033). Processing included merging tiled datasets, clipping to the study area, and enforcing strict pixel-wise alignment (snapping). Categorical variables were resampled using nearest-neighbor interpolation, while continuous variables were resampled using bilinear interpolation. This harmonization ensured consistency across all raster layers for subsequent modeling steps.

**Supplementary Methods 2. Regionalization framework**

To represent spatial heterogeneity, a data-driven regionalization approach was applied using spatially constrained clustering (Spatial 'K'luster Analysis by Tree Edge Removal - SKATER[1]). Considering an hexagonal with 20 km cells, regions were defined based on LULC proportions (2023) and transition rates (2018–2023), resulting in 24 spatially coherent regions (Fig. S1). This regionalization replaced administrative boundaries and allowed calibration and simulation to reflect localized land-use dynamics. All model calibration and simulation steps were conducted independently for each region.

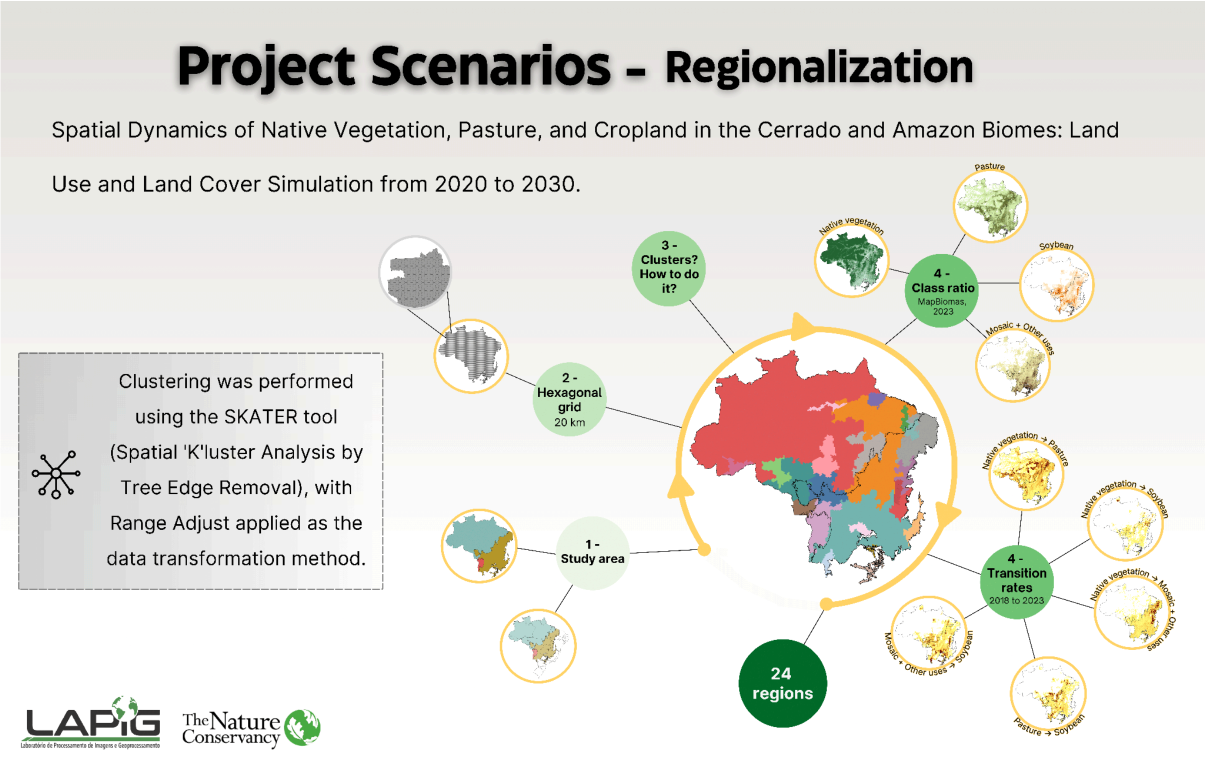

Figure S1. Regionalization of the Amazon, Cerrado, and Pantanal based on spatially constrained multivariate clustering (SKATER). Using a 20 km hexagonal grid and MapBiomas Collection 9 data, we generated 24 homogeneous regions defined by 2023 land-use proportions (Native Vegetation, Pasture, Soy, and Mosaic) and 2018–2023 transition rates.

**Supplementary Methods 3. Transition matrix and explanatory variables**

Model calibration began with the computation of a transition matrix based on the 2018 and 2023 LULC maps, quantifying the rate of change between all modeled transitions (Table S1). These rates provided the empirical basis for projections and were held constant during the 2023–2030 simulation period, while being estimated independently for each region.

A total of 39 explanatory variables were organized into four groups: (i) infrastructure and accessibility, (ii) land-use history, (iii) biophysical characteristics, and (iv) land tenure and institutional factors (Table S2).

To reduce redundancy, multicollinearity among variables was assessed using Cramér's V coefficient[2]. Variable pairs exceeding a threshold of 0.5 were evaluated (Table S3), and one variable from each pair was excluded based on ecological and methodological relevance[3]. This procedure ensured robust and interpretable probability surfaces.

Table S1. Land-use transitions considered in the modeling framework. The selected transitions, defined according to the objectives of this study, focused on the main land-use dynamics associated with native vegetation, pasture, and soybean, which are the dominant components of land-use change in Brazilian agricultural frontiers. By prioritizing these transitions, the analysis captured the primary pathways of land conversion and reallocation relevant to deforestation processes and agricultural intensification, while maintaining model parsimony. This approach is consistent with previous studies that highlight pasture expansion and cropland dynamics as the main drivers of land-use change in tropical regions[4,5].

| Transition | Description |
|---|---|
| Native vegetation to Pasture | Conversion of native vegetation to pastureland |
| Native vegetation to Other uses | Conversion of native vegetation to other temporary crops and mixed-use areas |
| Native vegetation to Soybean | Conversion of native vegetation to soybean cultivation |
| Pasture to Native vegetation | Conversion of pastureland to native vegetation |
| Pasture to Soybean | Conversion of pastureland to soybean cultivation |
| Other uses to Native vegetation | Conversion of temporary crops and mixed-use areas to native vegetation |
| Other uses to Soybean | Conversion of temporary crops and mixed-use areas to soybean cultivation |

Table S2. Summary of explanatory variables used in the LULC transition modeling. The variables are categorized into four thematic groups: (i) Infrastructure and Accessibility; (ii) Land Use and Land Cover; (iii) Biophysical and Natural Factors; and (iv) Land Tenure and Institutional Factors.

| **Category** | **Variable** | **Description** | **Year** | **Source** |
| --- | --- | --- | --- | --- |
| Anthropogenic | Permanent Preservation Area (APP) | Vector split into APP and Legal Reserve; rasterized, reprojected, resampled, and aligned. | — | SICAR |
| Anthropogenic | Legal Reserve | Raster derived from SICAR; reprojected, resampled, and aligned. | — | SICAR |
| Anthropogenic | Brazilian Forest Code | Overlay of biome, states, and vegetation; classified into legal requirement classes; resampled to 30 m. | 2023 | IBGE |

| | | | | |
|---|---|---|---|---|
| Anthropogenic | Indigenous land | Vector reprojected, rasterized, resampled, and clipped. | 2025 | FUNAI |
| Anthropogenic | Strictly Protected Area | Protected areas categorized and rasterized; resampled to 30 m. | 2025 | MMA |
| Anthropogenic | Sustainable Protected Area | Protected areas categorized and rasterized; resampled to 30 m. | 2025 | MMA |
| Anthropogenic | Territorial unit | Classified into public, private, settlements, and *quilombola*; rasterized and aligned. | 2022, 2025 | MMA, INCRA |
| Anthropogenic | Property Size | Reclassified into four classes; rasterized, reprojected, and aligned. | — | INCRA |

| | | | | |
|---|---|---|---|---|
| Anthropogenic | Population density | Merged vector data converted to raster; reprojected and aligned. | 2010 | IBGE |
| Anthropogenic | Soybean Suitability | Filtered and reclassified into suitability classes; resampled to 30 m. | — | IBGE |
| Anthropogenic | Livestock Intensification | Scenario-based layer representing intensification potential of pasture systems. | — | TNC |
| Anthropogenic | Distance to Paved Federal and State Roads | Euclidean distance from paved road network. | 2021–2022 | DNIT |
| Anthropogenic | Unpaved Roads | Euclidean distance from unpaved road network. | 2023 | IBGE |
| Anthropogenic | Distance to Railways | Euclidean distance from railway network. | 2020 | DNIT |

| | | | | |
|---|---|---|---|---|
| Anthropogenic | Distance to Railway Terminals | Euclidean distance from railway terminals. | — | Various |
| Anthropogenic | Distance to Slaughterhouses | Euclidean distance from slaughterhouse facilities. | 2021 | ABIEC et al. |
| Anthropogenic | Distance to Warehouses | Euclidean distance from storage infrastructure. | 2024 | MapBiomas |
| Anthropogenic | Distance to River Ports | Euclidean distance from river ports. | — | Various |
| Anthropogenic | Distance to Navigable Waterways | Euclidean distance from navigable rivers. | 2022 | DNIT |
| Anthropogenic | Distance to Illegal Mining Sites | Euclidean distance from illegal mining areas. | — | Various |
| Anthropogenic | Distance to Rivers | Euclidean distance from drainage network. | 2017 | SNIRH |
| Anthropogenic | Deforestation Age | Year of vegetation conversion derived from | 2024 | MapBiomas |

| | | | | |
|---|---|---|---|---|
| | | MapBiomas historical series. | | |
| Anthropogenic | 2018 Land Cover | Land use and land cover classification. | 2018 | MapBiomas |
| Anthropogenic | 2023 Land Cover | Land use and land cover classification. | 2023 | MapBiomas |
| Anthropogenic | 2018 Pasture Age | Derived from MapBiomas pasture age product. | 2018 | MapBiomas |
| Anthropogenic | 2023 Pasture Age | Derived from MapBiomas pasture age product. | 2023 | MapBiomas |
| Anthropogenic | 2018 Pasture Vigor | Derived from EVI-based pasture vigor classification. | 2018 | LAPIG/MapBiomas |
| Anthropogenic | 2023 Pasture Vigor | Derived from EVI-based pasture vigor classification. | 2023 | LAPIG/MapBiomas |

| | | | | |
|---|---|---|---|---|
| Natural | Precipitation | Climatic layer reprojected and resampled to 30 m. | 2021 | CHELSA |
| Natural | Relief | Derived from ALOS PALSAR DEM. | 2015 | ASF Data Search |
| Natural | Slope | Derived from NASA DEM. | 2020 | NASA |
| Natural | Vegetation physiognomy | Classification of vegetation types across the study area. | — | IBGE |
| Natural | 2018 Median Vegetation Height | Raster representing vegetation height. | 2018 | LAPIG/GPW |
| Natural | 2022 Median Vegetation Height | Raster representing vegetation height. | 2022 | LAPIG/GPW |

Table S3. Correlation screening outcomes (Cramér's V > 0.5) for region 3, for example purposes. The material for this step is extensive and is available in the GitHub repository. The numbers below refer to the MapBiomas classes.

| Transition | Variable 1 | Variable 2 | Cramér's V | Variable retained | Rationale |
|---|---|---|---|---|---|
| 1-15 | Deforestation | Pasture Age | 0.6237 | Deforestation | Primary conversion proxy |
| 1-21 | Deforestation | Pasture Age | 0.6121 | Deforestation | Primary conversion proxy |
| 1-39 | Deforestation | Pasture Age | 0.6162 | Deforestation | Primary conversion proxy |
| 15-1 | Deforestation | Pasture Age | 0.6133 | Pasture Age | Legacy of prior use |
| 15-39 | Deforestation | Pasture Age | 0.6133 | Pasture Age | Legacy of prior use |
| 21-1 | Deforestation | Pasture Age | 0.5715 | Deforestation | Primary disturbance legacy |
| 21-39 | Deforestation | Pasture Age | 0.6124 | Deforestation | Primary disturbance legacy |

**Supplementary Methods 4. Weights of Evidence calibration and spatial allocation**

Spatial allocation of land-use transitions was performed using the Weights of Evidence (WoE) method[2], which estimates the statistical association between explanatory variables and observed transitions using a Bayesian framework. For each variable, distance-based binning was applied to maximize contrast between transition and non-transition areas. WoE values were then used as partial probability modifiers within the Dinamica EGO allocation process (Table S4).

Despite low statistical significance during calibration, legal constraints (LRs, PPAs, Strictly Protected Areas, and Indigenous Lands) were incorporated as normative weights to represent institutional restrictions. These weights were not empirically derived but imposed to simulate policy-relevant scenarios.

Spatial allocation was controlled using the Patcher and Expander functions in Dinamica EGO, which regulate patch size distribution and the balance between expansion and new patch formation[6]. Parameterization was based on landscape metrics such as mean patch size, variance, and isometry[7].

Table S4. Summary of Weights of Evidence (WoE) at the variable level, reporting the number of regions in which each variable was retained after correlation tests, the number of transitions in which it was used, and the mean WoE across region–transition combinations. WoE values indicate the strength and direction of association between explanatory variables and land-use transitions. The complete set of region-specific WoE tables, including all distance bins, is available in the project repository (https://github.com/lapig-ufg/Scenarios-project).

| **Variable** | **Regions used** | **Transitions involved** | **Mean WoE** |
|---|---|---|---|
| Pasture Vigor | 24 | 7 | -0.016 |
| Deforestation Age | 24 | 7 | -0.178 |

| | | | |
|---|---|---|---|
| Distance to Navigable Waterways | 24 | 7 | -0.204 |
| Distance to Paved Federal and State Roads | 24 | 7 | -0.253 |
| Distance to Rivers | 24 | 7 | -0.126 |
| Distance to Slaughterhouses | 24 | 7 | -0.18 |
| Distance to Warehouses | 24 | 7 | -0.391 |
| Population density | 24 | 7 | -0.155 |
| Precipitation | 24 | 7 | -0.209 |
| Slope | 24 | 7 | -0.418 |
| Soybean Suitability | 24 | 7 | -0.176 |
| Distance to Pasture | 24 | 5 | -0.535 |
| Distance to Native vegetation | 24 | 4 | -0.644 |
| Distance to River Ports | 23 | 7 | -0.271 |
| Property Size | 23 | 7 | 0.037 |
| Relief | 23 | 7 | -0.22 |
| Territorial unit | 23 | 7 | -0.225 |
| Unpaved Roads | 23 | 7 | -0.262 |
| Vegetation physiognomy | 23 | 7 | -0.423 |
| Distance to Other uses | 23 | 5 | -0.839 |

| | | | |
|---|---|---|---|
| Distance to Soybean | 22 | 7 | -0.608 |
| Pasture Age | 20 | 7 | 0.029 |
| Permanent Preservation Area (APP) | 19 | 7 | -0.26 |
| Distance to Illegal Mining Sites | 18 | 7 | -0.395 |
| Distance to Railways | 17 | 7 | -0.237 |
| Distance to Railway Terminals | 16 | 7 | -0.298 |
| Brazilian Forest Code | 14 | 7 | -0.049 |
| Legal Reserve | 2 | 7 | -0.25 |
| Strictly Protected Area | 2 | 3 | 0.0 |
| Sustainable Protected Area | 2 | 3 | 0.0 |
| Indigenous land | 1 | 2 | 0.0 |

**Supplementary Methods 5. Scenario parameterization**

Three scenarios were defined to represent distinct governance and intensification pathways (Table S5):

- BAU (Business as Usual): maintains observed transition rates and baseline normative weights (PPAs and protected areas = 2; LRs = 1).
- CG (Collaborative Governance): removes native vegetation-to-pasture transitions and increases normative weights (PPAs and protected areas = 4; LRs = 2).

- IG (Integrated Governance): retains CG restrictions and incorporates a livestock intensification suitability layer derived from the TNC degraded pasture framework, based on field data and expert curation aligned with PNCPD (i.e. national program on degraded pasture conversion).

In IG, only the WoE value associated with pasture-to-native vegetation transitions (−1.5) was maintained, thereby discouraging restoration-related transitions and directing production toward already established pasture areas.

Table S5. Scenario parameterization, with the differences used in the modeling approach.

| | **Scenarios** | | |
|---|---|---|---|
| **Component** | **BAU** | **CG** | **IG** |
| Native vegetation to Pasture | Allowed | Blocked | Blocked |
| Permanent Preservation Areas (APP) | 2 | 4 | 4 |
| Legal Reserves (RL) | 1 | 2 | 2 |
| Indigenous lands | 2 | 4 | 4 |
| Strictly Protected Areas | 2 | 4 | 4 |
| Intensification layer | no | no | yes (WoE = −1.5) |

**Supplementary Methods 6. Validation procedures**

Model validation compared simulated and observed 2023 maps using two similarity metrics: exponential decay and constant decay. Similarity was assessed using moving windows ranging from 3×3 to11×11 pixels.

The exponential decay metric assigns partial similarity to nearby transitions, while the constant decay metric considers transitions correct if they occur within the defined window.

The model was considered valid if:

- exponential similarity ≥ 0.5, or
- constant similarity ≥ 0.4.

Results are reported for the 11×11 window (Table S5). When thresholds were not met, Patcher and Expander parameters were recalibrated.

Table S5. Summary of model validation results across regions, reporting the mean similarity values for exponential and constant decay functions using a 11×11 moving window. For each region, values represent the average between minimum and maximum similarity scores obtained during validation.

| **Region** | **Exponential decay similarity** | **Constant decay similarity** |
|---|---|---|
| 1 | 0.27 | 0.40 |
| 2 | 0.40 | 0.61 |
| 3 | 0.39 | 0.67 |
| 4 | 0.36 | 0.53 |
| 5 | 0.43 | 0.62 |
| 6 | 0.30 | 0.40 |
| 7 | 0.44 | 0.64 |
| 8 | 0.52 | 0.71 |

| | | |
|---|---|---|
| 9 | 0.52 | 0.70 |
| 10 | 0.29 | 0.48 |
| 11 | 0.47 | 0.62 |
| 12 | 0.65 | 0.79 |
| 13 | 0.40 | 0.70 |
| 14 | 0.44 | 0.65 |
| 15 | 0.60 | 0.77 |
| 16 | 0.58 | 0.78 |
| 17 | 0.45 | 0.58 |
| 18 | 0.36 | 0.53 |
| 19 | 0.43 | 0.64 |
| 20 | 0.29 | 0.47 |
| 21 | 0.61 | 0.74 |
| 22 | 0.66 | 0.83 |
| 23 | 0.43 | 0.63 |
| 24 | 0.33 | 0.58 |

**Supplementary Methods 7. Post-simulation calculations**

### *Avoided deforestation*

Avoided deforestation (AD) was calculated as the difference in native vegetation loss between the BAU scenario and each alternative governance scenario (CG and IG) over the 2023–2030 period. Native vegetation loss was first quantified for the BAU scenario as the total area converted from native vegetation to other land-use classes between the initial (2023) and final (2030) simulated maps. The same procedure was applied to each alternative scenario.

Avoided deforestation was then calculated as:

$$\mathbf{AD = Loss\_BAU - Loss\_scenario}$$

where *Loss_BAU* represents native vegetation loss under the BAU scenario, and *Loss_scenario* represents loss under CG or IG.

All calculations were performed at 30 m spatial resolution and aggregated to million hectares (Mha). Spatially explicit results were further summarized using a hexagonal grid (20 $km^2$ cells), where the proportion of native vegetation remaining (i.e., not converted) was calculated for each cell.

#### *Cropland and pasture reallocation analysis*

Cropland and pasture reallocation were assessed by comparing the 2023 MapBiomas land-use map with each simulated 2030 map on a pixel-by-pixel basis. Newly converted soybean areas were identified as pixels classified as soybean in 2030 that belonged to a different land-use class in 2023. For each of these pixels, the land-use class of origin was recorded, allowing the identification of transition flows from native vegetation, pasture, and other land-use classes (Table S6).

Transition flows were quantified both as total area (million hectares, Mha) and as a proportion of total soybean expansion under each scenario.

All calculations were performed at 30 m spatial resolution, and area estimates were derived from pixel counts converted to hectares.

Table S6: Soybean expansion origin by scenario.

| Scenario | Soybean Expansion (ha) | From Native (ha) | From Pasture (ha) | From Other Uses (ha) | From Native (%) | From Pasture (%) | From Other (%) |
|---|---|---|---|---|---|---|---|
| Business as Usual | 6,262,152.84 | 555,703.11 | 755,264.79 | 4,951,184.94 | 8.87% | 12.06% | 79.07% |
| Collaborative Governance | 6,226,582.41 | 471,179.97 | 712,779.84 | 5,042,622.6 | 7.57% | 11.45% | 80.99% |
| Integrated Governance | 6,226,789.05 | 473,444.01 | 713,214.81 | 5,040,130.23 | 7.60% | 11.45% | 80.94% |

#### *Livestock intensification requirement*

Livestock intensification requirements were estimated as the ratio between total pasture area under the BAU scenario and that under each alternative scenario (CG and IG). This ratio was applied to a baseline stocking rate of 1.00 AU/ha to estimate the effective stocking rate required to maintain equivalent livestock production under reduced pasture area.

**Supplementary Methods 8. Expanded LLI note**

The Landscape Line Intercept (LLI) was computed for each hexagonal grid cell (20 km²) as the combination of two components: (i) native vegetation cover and (ii) edge continuity. Native vegetation cover was calculated as the proportion of pixels classified as native vegetation within each hexagonal cell:

$$\mathbf{Cover = Nveg / Ntotal}$$

where *Nveg* is the number of pixels classified as native vegetation and *Ntotal* is the total number of valid pixels within the cell.

Edge continuity was calculated as the proportion of perimeter-intersecting pixels classified as native vegetation:

$$\mathbf{Continuity = Nedge_{veg} / Nedge_{total}}$$

where $Nedg_{eveg}$ is the number of pixels intersecting the cell boundary classified as native vegetation, and $Nedge_{total}$ is the total number of pixels intersecting the boundary.

Both components were derived from binary native vegetation rasters (value 1 = native vegetation; value 0 = non-vegetation) using zonal statistics. The LLI formulation integrates habitat amount and boundary continuity within each spatial unit, capturing structural properties associated with landscape fragmentation, edge effects, and ecological connectivity.

Table S7. Hexagon-class frequencies for landscape structure (BAU 2025 vs BAU 2030)

| Native vegetation (%) | Continuity (%) | BAU 2025 (%) | BAU 2030 (%) | Difference |
|---|---|---|---|---|
| 80–100 | 0–20 | 0.0 | 0.0 | 0.0 |
| 80–100 | 20–40 | 0.0 | 0.0 | 0.0 |
| 80–100 | 40–60 | 0.0 | 0.0 | 0.0 |
| 80–100 | 60–80 | 2.1 | 2.2 | +0.1 |
| 80–100 | 80–100 | 54.0 | 53.1 | −0.9 |

**Limitations**

Several limitations of this study warrant acknowledgment. The weights of evidence assigned to protected areas were not derived from empirical calibration; although we tested alternative weight configurations, no established benchmarks exist in the literature for quantifying institutional enforcement intensity in spatially explicit land-use models, and the values adopted represent theoretically grounded scenario parameters rather than measured compliance rates. The Landscape Line Intercept (LLI) was developed specifically for this study and has not been externally validated against established landscape ecology metrics, representing an avenue for future methodological refinement. The SICAR and SIGEF datasets, despite known coverage limitations, underwent systematic preprocessing and expert curation by TNC specialists prior to integration, partially mitigating data quality concerns. The livestock intensification suitability layer was derived from a single data source; however, this layer was constructed from field data and expert curation by TNC Brazil and represents the most comprehensive spatially explicit information currently available for the study area. Finally, spatial validation was performed across the full range of moving window sizes from 3×3 to 11×11 pixels; results reported here correspond to the 11×11 window, which provided the most balanced representation of neighborhood-level land-use dynamics at the spatial resolution of the input data

**Supplementary Results**

### Supplementary Results 1. Full three-scenario outputs

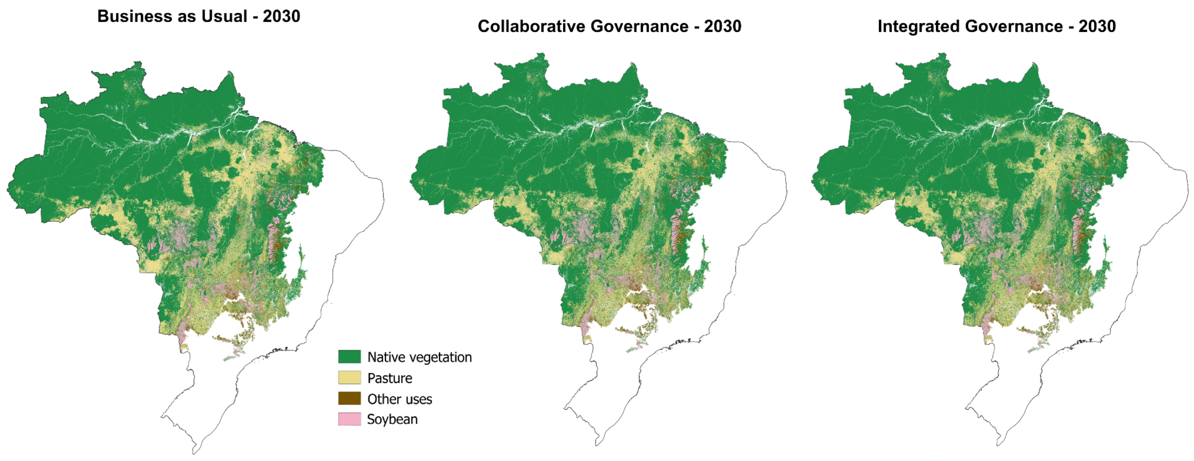


Figure S2. Land-use patterns in 2030 under different governance regimes. Simulated land-use maps for 2030 under Business as Usual (BAU), Collaborative Governance (CG), and Integrated Governance (IG), showing the spatial distribution of native vegetation, pasture, soybean, and other land-use classes.

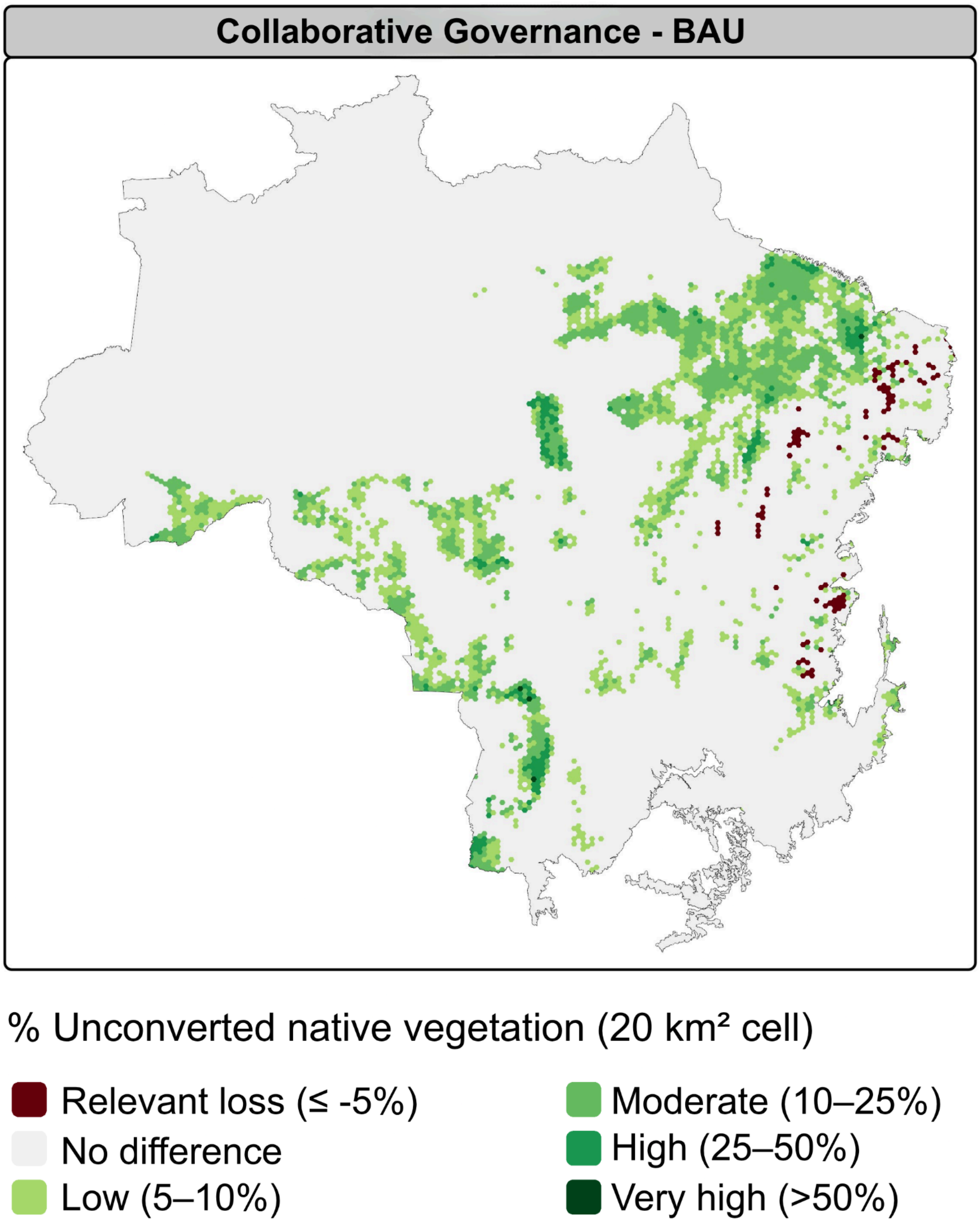


Figure S3. Avoided native vegetation loss under Collaborative Governance relative to BAU (2023-2030). Spatial distribution of avoided native vegetation loss under the Collaborative Governance (CG) scenario relative to Business as Usual (BAU), expressed as the percentage of unconverted native vegetation within each 20 km² hexagonal cell. Color classes indicate increasing levels of avoided loss.

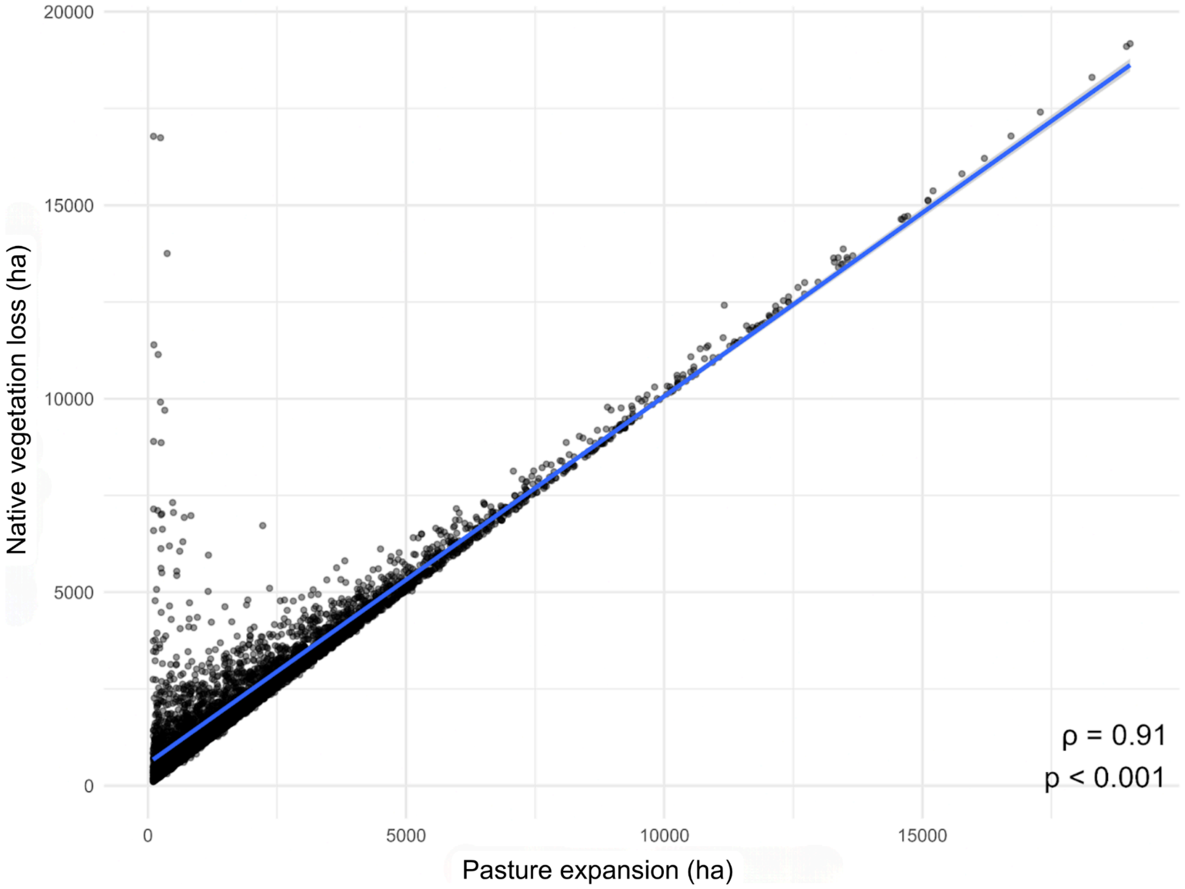


Figure S4. Relationship between pasture expansion (ha) and native vegetation loss (ha) under the BAU scenario, at the analysis unit level. Each point represents a spatial unit, and the solid line indicates the fitted linear trend. A strong positive relationship is observed (Spearman $\rho = 0.91$; $p < 0.001$).

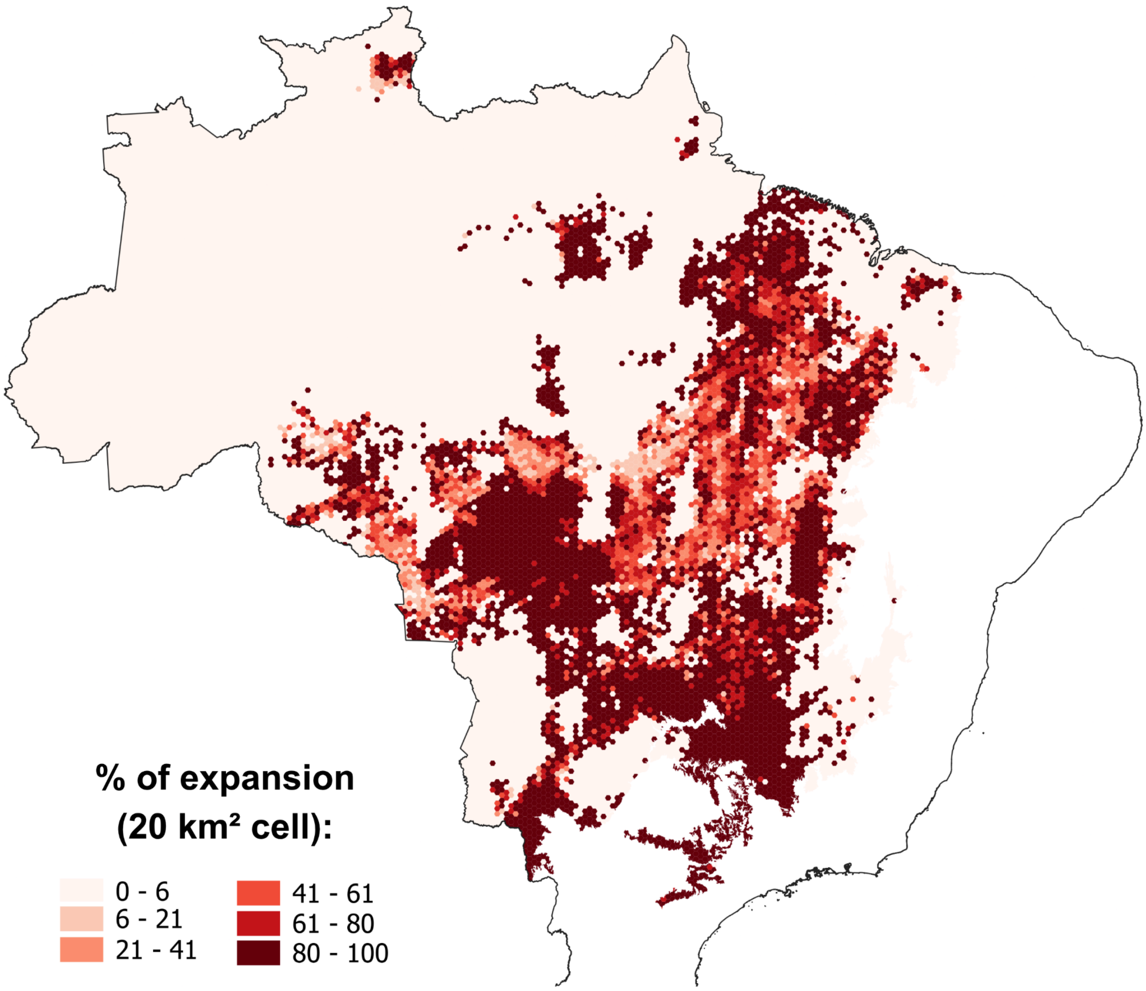


Figure S5. Soybean expansion from other land-use classes. Proportion of soybean expansion (2023–2030) originating from other land-use classes across Integrated Governance regime.

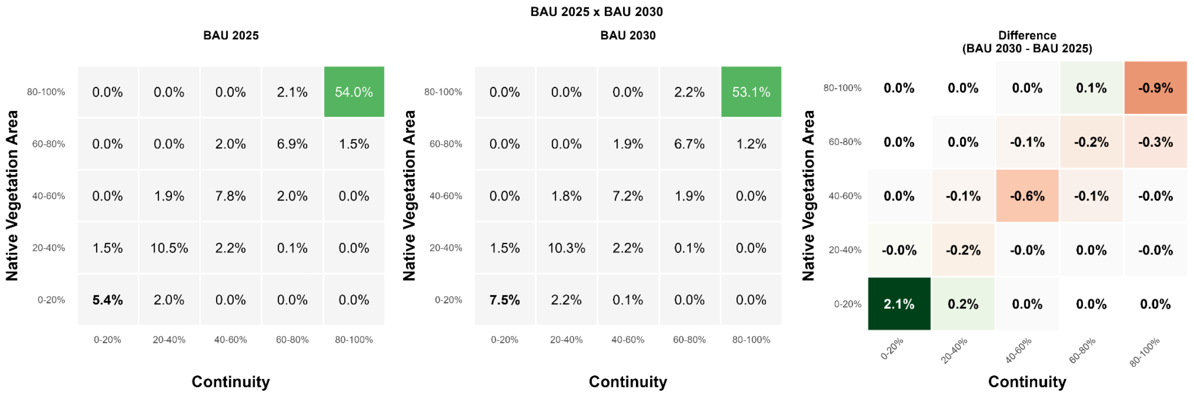


Figure S6. Changes in landscape structure under BAU between 2025 and 2030. (A) Relationship between native vegetation area (%) and landscape continuity (%) under the Business as Usual (BAU) scenario in 2025, with the corresponding distribution of hexagons across vegetation cover and continuity classes. (B) Same relationship under BAU in 2030. (C) Difference between 2030 and 2025 under BAU, expressed as changes in the proportion of hexagons across vegetation cover and continuity classes.

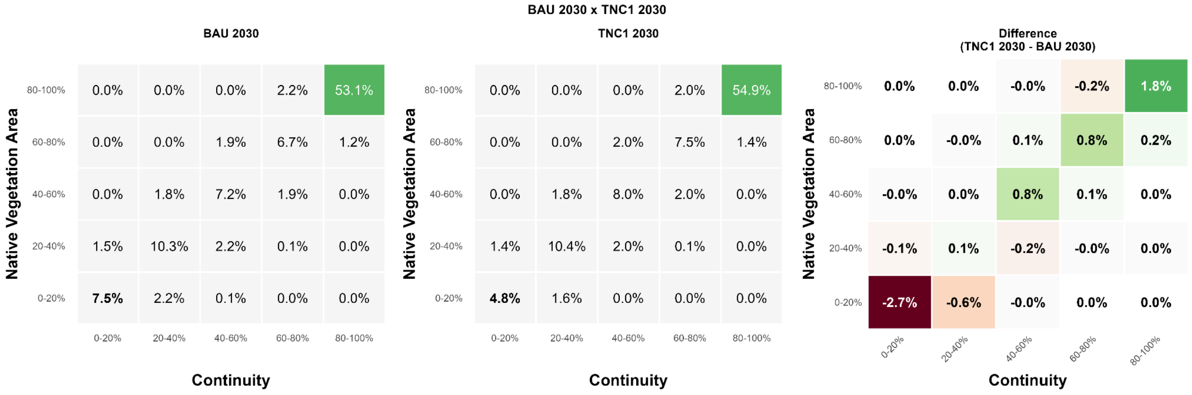


Figure S7. Changes in landscape structure under Collaborative Governance relative to BAU. (A) Relationship between native vegetation area (%) and landscape continuity (%) under the Business as Usual (BAU) scenario in 2030, with the corresponding distribution of hexagons across vegetation cover and continuity classes. (B) Same relationship under the Collaborative Governance (CG) scenario in 2030. (C) Difference between CG and BAU in 2030, expressed as changes in the proportion of hexagons across vegetation cover and continuity classes.
